# Knockout of cardiolipin synthase disrupts postnatal cardiac development by inhibiting the maturation of mitochondrial cristae

**DOI:** 10.1101/2023.03.09.531996

**Authors:** Mindong Ren, Yang Xu, Colin K. L. Phoon, Hediye Erdjument-Bromage, Thomas A. Neubert, Michael Schlame

## Abstract

**Background:** Cardiomyocyte maturation requires a massive increase in respiratory enzymes and their assembly into long-lived complexes of oxidative phosphorylation (OXPHOS). The molecular mechanisms underlying the maturation of cardiac mitochondria have not been established.

**Methods:** To determine whether the mitochondria-specific lipid cardiolipin is involved in cardiac maturation, we created a cardiomyocyte-restricted knockout (KO) of cardiolipin synthase (*Crls1*) in mice and studied the postnatal development of the heart. We also measured the turnover rates of proteins and lipids in cardiolipin-deficient flight muscle from Drosophila, a tissue that has mitochondria with high OXPHOS activity like the heart.

**Results:** *Crls1KO* mice survived the prenatal period but failed to accumulate OXPHOS proteins during postnatal maturation and succumbed to heart failure at the age of 2 weeks. Turnover measurements showed that the exceptionally long half-life of OXPHOS proteins is critically dependent on cardiolipin.

**Conclusions:** Cardiolipin is essential for the postnatal maturation of cardiomyocytes because it allows mitochondrial cristae to accumulate OXPHOS proteins to a high concentration and to shield them from degradation.

## INTRODUCTION

Cardiolipin (CL) is a mitochondrial phospholipid discovered in and named after the lipid extract of bovine heart (Pangborn, 1942). The high abundance of CL in heart suggests that it is significant for cardiac function. Indeed, CL deficiency is associated with cardiomyopathy in patients with Barth syndrome (Schlame et al, 2003; Vreken et al, 2000), in patients with CL synthase (*Crls1*) mutations (Lee et al, 2022), and in diabetic mice (Han et al, 2005; 2007).

There are multiple reasons why CL might be important in the heart. CL is essential for energy efficiency, which has been attributed to its interaction with OXPHOS proteins (Claypool et al, 2008; Xu et al 2021). Energy efficiency, and therefore CL, is likely to be critical in an energy-demanding organ like heart (Bertero et al, 2020). Apart from that, CL might play a role in heart development. Cardiomyocytes differentiate through a tightly scripted program that prepares intracellular organelles for the task of perpetual contraction (Karbassi et al, 2020). In mature cardiomyocytes, mitochondria are unique in their extreme state of fusion, high crista density, and spatial alignment with myofibrils (Hollander et al, 2014), which all may require specific lipids, such as CL. Heart mitochondria are built to last extended periods of time (Fornasiero et al, 2018; Kim et al, 2012). In particular the inner membrane of heart mitochondria, where CL is localized, harbors proteins of remarkable longevity, lasting up to several months (Bomba-Warczak et al, 2021). Since CL itself has a very long half-life (Landriscina et al, 1976; Wahjudi et al, 2011; Xu et al, 2016), the question arises as to whether CL is involved in conferring longevity to the inner membrane. In short, the specific function of CL and its mechanism of action both in heart development and in heart function, remain important unresolved issues.

To gain further insight into the role of CL, a knockout (KO) mouse model is needed. However, constitutive KO of CL synthesis in mice results in early embryonic lethality (Kasahara et al, 2020; Zhang et al, 2011). Tissue-specific KOs of enzymes involved in CL synthesis, specifically *Crls1* and *Ptpmt1*, have been accomplished in brown adipocytes (Sustarsic et al, 2018), neurons (Kasahara et al, 2020), and T-cells (Corrado et al, 2020). Those mouse strains have confirmed previously known effects of CL deficiency and uncovered novel tissue-specific functions in thermogenesis control and T cell differentiation, but they have not been able to address the specific function of CL in the heart.

To close this gap, a cardiomyocyte-restricted KO of *Ptpmt1* has been created (Chen et al, 2021). This has induced CL deficiency in embryonic mouse hearts, but again prenatal lethality has limited the application of this model. Another CL deficiency model has been established by cardiomyocyte-restricted KO of tafazzin (Wang et al, 2020), the enzyme that remodels the acyl chains of CL after de novo synthesis (Schlame & Xu, 2020). However, in this instance the effects of low CL are confounded by an abnormal species composition and by the accumulation of monolyso-CL, a variant of CL with 3 acyl chains.

Here we have created a new mouse model with cardiomyocyte-restricted KO of *Crls1*, in which the myocardial concentration of CL is severely reduced. Mice were born with normal cardiac function but soon developed cardiomyopathy and died within 2 weeks. This model provides a unique opportunity to study the mechanism of action of CL, specifically during the maturation of cardiomyocytes in the neonatal period.

## METHODS

### Animals

All protocols were approved by the Institutional Animal Care and Use Committee of the NYU School of Medicine and conform to the Guide for the Care and Use of Laboratory Animals published by the National Institutes of Health (NIH). Mice were housed under temperature-controlled conditions under a 12-hour light/dark cycle with free access to drinking water and food.

Mice with floxed cardiolipin synthase gene (*Crls1*^flox/flox^) were generated by crossing the conditional ready strain (C57BL/6N-A^tm1Brd^ Crls1^tm1a(EUCOMM)Wtsi^/WtsiOulu) from the European Mouse Mutant Archive with the FLPo deleter strain (B6.129S4-*Gt(ROSA)26Sor^tm2(FLP*)Sor^*/J, Strain # 012930) from The Jackson Laboratory to remove the reporter-tagged insertion. Crls1^flox/flox^ mice were crossed with the *αMyHC-cre* mouse (B6.FVB-Tg[Myh6-cre]2182Mds/J, Strain # 011038) from The Jackson Laboratory, to generate *αMyHC-cre;Crls1^flox/+^* mice. Cardiomyocyte-specific *Crls1*-KO mice (αMyHC-cre; Crls1^flox/flox^) were generated by crossing Crls1^flox/flox^ mice with *αMyHC-cre;Crls1^flox/+^* mice. Genotyping was performed by polymerase chain reaction in tail clips or in yolk sacs of embryos.

Hearts were harvested from mice euthanized by CO2 narcosis/asphyxiation. Hearts were excised through a vertical anterior thoracotomy, trimmed from attached noncardiac tissue, rinsed in cold phosphate-buffered saline, and either homogenized in water or processed for the purification of cardiomyocytes. In order to purify cardiomyocytes, hearts were dissociated into single-cell suspensions using the Neonatal Heart Dissociation kit from Miltenyi Biotec, which applies a combination of enzymatic degradation and mechanical disruption. Non-cardiomyocytes were removed from the cell suspension by affinity binding to monoclonal antibodies. For that purpose, we used the Neonatal Cardiomyocyte Isolation kit from Miltenyi Biotec. Heart and cardiomyocyte samples were stored at −80 °C. The protein concentration was determined by the method of Lowry (Lowry et al, 1951).

Drosophila strains were maintained on standard fly food containing yeast, cornmeal, and molasses in 3-inch culture vials at 24 °C. The strain w^1118^; PBac{PB} CG4774^c01874^/TM6B, Tb1, with a transposon insertion in the coding region of the last exon of the cardiolipin synthase gene (*ΔCrls1*), was obtained from the Bloomington Drosophila Stock Center (No. 10741). The tafazzin mutant (*ΔTaz*) was created in our laboratory (Xu et al, 2006). To avoid confounding effects, we re-derived all strains in identical genetic backgrounds (Xu et al, 2021).

### Echocardiography

Transthoracic echocardiography was performed on anesthetized mice as previously described (Phoon et al, 2012; Phoon & Turnbull, 2016). In brief, mice aged 5 days and older were anesthetized with isoflurane via nose cone (induction: 2.5% isoflurane; maintenance: 1% isoflurane at a flow rate of 1l/min) with strict thermoregulation (37±0.5°C) to optimize physiological conditions and reduce hemodynamic variability. Because depilation of the chest was not necessary in younger mice, they were gently held and quickly scanned (<1 minute) without anesthesia. Two-dimensional echocardiography of the parasternal short axis view was performed using the Vevo 770 (Visual Sonics, Toronto, ON, Canada) with the RMV-704, 40 MHz center frequency transducer (age>4 days) or the RMV-711, 55 MHz center frequency transducer (age<4 days). To minimize bias, we performed all echocardiographic measurements offline while blinded to the genotype.

### Microscopy

Paraffin sections of 6 μm thickness were prepared for histological evaluations of the hearts. The sections were stained with hematoxylin and eosin. For EM, tissue samples (<3 mm diameter) were fixed in glutaraldehyde and osmium tetraoxide. Fixed samples were stained with uranyl acetate. The buffer was gradually exchanged with ethanol, followed by resin exchange and heat polymerization. Sections of 50–100 nm were cut with a Leica Ultracut UCT microtome and collected on EM grids. Sections were stained with uranyl acetate and Sato Lead stains and imaged with a Philips CM12 transmission electron microscope. Random images were collected at different magnifications. We determined the areas of mitochondrial cross sections at a magnification of 2,500. Crista densities were determined at a magnification of 5,000. Crista density was defined as the number of cristae that intersect a line drawn across the mitochondrion.

### CRLS1 activity assay

To determine CRLS1 activity, we measured the formation of a non-natural CL species (CL14:0/14:0/18:1/18:1) during incubation of heart tissue in the assay medium. Heart homogenate, corresponding to 70 μg protein, was incubated for 45 minutes at 37°C in 0.1 mL buffer containing 10 mM Tris (pH 8.0), 5 mM CoCl2, 0.1 mM dimyristoylphosphatidylglycerol (PG14:0/14:0), and 0.1 mM CDP-dioleoylglycerol (CDPDG18:1/18:1). Reactions were stopped by the addition of 2 mL methanol and 1 mL chloroform. Cardiolipin standard mix I (Avanti Polar Lipids) was added as internal standard, containing 101 pmol CL14:1/14:1/14:1/15:1, 92.2 pmol CL15:0/15:0/15:0/16:1, 84.7 pmol CL14:1/22:1/22:1/22:1, and 77.1 pmol CL14:1/24:1/24:1/24:1. Lipids were extracted (Bligh & Dyer, 1959), dried under nitrogen, and re-suspended into 0.1 mL chloroform/methanol 1:1. An aliquot of 5 μL was applied to LC-MS/MS analysis as described below. To determine the amount of CL14:0/14:0/18:1/18:1 formed during the incubation, signal intensities of CL14:0/14:0/18:1/18:1 and of 2 CL standards (CL14:1/14:1/14:1/15:1, CL15:0/15:0/15:0/16:1) were quantified in the Qual Browser of Xcalibur (Thermo Fisher Scientific, West Palm Beach, FL 33407, Version 4.1.50). We have shown that the concentration of CL14:0/14:0/18:1/18:1 rises in linear relation to the incubation time and to the amount of heart homogenate. CRLS1 activities were expressed as fmol CL14:0/14:0/18:1/18:1 formed per minute and μg protein.

### Lipidomics

Samples, containing about 1 mg cardiac protein, were extracted into chloroform/methanol as described (Bligh & Dyer, 1959). To this end, samples were homogenized in 0.2 mL water, suspended in methanol/chloroform (2:1), and incubated at 37°C for 30 minutes to denature proteins. A mixture of internal standards was added consisting of Cardiolipin standard mix I (303 pmol CL14:1/14:1/14:1/15:1, 277 pmol CL15:0/15:0/15:0/16:1, 254 pmol CL14:1/22:1/22:1/22:1, 231 pmol CL14:1/24:1/24:1/24:1) and Mouse SPLASH (625 pmol PC15:0/18:1d7, 44 pmol PE15:0/18:1d7, 125 pmol PS15:0/18:1d7, 31 pmol PG15:0/18:1d7, 125 pmol PI15:0/18:1d7, 62.5 pmol PA15:0/18:1d7, 281 pmol LPC18:1d7, 12.5 pmol LPE18:1d7, 1562 pmol cholesterol ester18:1d7, 125 pmol plasmenyl-PC18:0/18:1d9, 94 pmol DG15:0/18:1d7, 219 pmol TG15:0/18:1d7/15:0, 125 pmol SMd18:1/18:1d9, 31 pmol plasmenyl-PE18:0/18:1d9). Standards were obtained from Avanti Polar Lipids. Chloroform and water were added, the samples were vortexed, and phase separation was achieved by centrifugation. The lower phase was collected, dried under nitrogen, and re-dissolved in 0.2 ml chloroform/methanol (1:1). Lipids were analyzed by LC-ESI-MS/MS on a QExactive HF-X instrument coupled directly to a Vanquish UHPLC (Thermo Fisher Scientific, Waltham, MA, USA). An aliquot of 5 μl was injected into a Restek Ultra C18 reversed-phase column (Restek Corporation, Bellefonte, PA, USA; 100×2.1 mm; particle size 3 μm) that was kept at a temperature of 50°C. Chromatography was performed with solvents A and B at a flow rate of 0.15 mL/min. Solvent A contained 600 ml acetonitrile, 399 ml water, 1 ml formic acid, and 0.631 g ammonium formate. Solvent B contained 900 ml 2-propanol, 99 ml acetonitrile, 1 ml formic acid, and 0.631 g ammonium formate. The chromatographic run time was 40 minutes, changing the proportion of solvent B in a non-linear gradient from 30 to 35% (0-2 minutes), from 35 to 67% (2-5 minutes), from 67 to 83% (5-8 minutes), from 83 to 91% (8-11 minutes), from 91 to 95% (11-14 minutes), from 95 to 97% (14-17 minutes), from 97 to 98% (17-20 minutes), from 98 to 100% (20-25 minutes), and from 100 to 30% (25-26 minutes). For the remainder of the run time the proportion of solvent B stayed at 30% (26-40 minutes). To analyze acyl-carnitines, cholesterol esters, and glycerides, the mass spectrometer was operated in positive ion mode. For all other lipids, it was operated in negative ion mode. The spray voltage was set to 4 kV and the capillary temperature was set to 35O°C. MS1 scans were acquired in profile mode at a resolution of 120,000, an AGC target of 1e6, a maximal injection time of 65 ms, and a scan range of 200-2000 m/z. MS2 scans were acquired in profile mode at a resolution of 30,000, an AGC target of 3e6, a maximal injection time of 75 ms, a loop count of 7, and an isolation window of 1.7 m/z. The normalized collision energy was set to 30 and the dynamic exclusion time to 31 s. For lipid identification and quantitation, data were analyzed by the software LipidSearch 5.0 (Thermo Fisher Scientific, Waltham, MA, USA). The general database was searched with a precursor tolerance of 4 ppm, a product tolerance of 10 ppm, and an intensity threshold of 1.0%.

### Proteomics

Protein abundances in heart samples were determined by label-free relative quantitative proteomics (MaxLFQ) using data-independent acquisition (MaxDIA) (Sinitcyn et al, 2021). Proteins were purified and concentrated by a brief SDS-PAGE run designed to focus all proteins into a single band of about 1 cm x 0.5 cm. Gels were washed 3 times in double-distilled water for 15 minutes each. Proteins were visualized and fixed by staining with EZ-Run Protein staining solution (ThermoFisher Scientific, USA). The stained protein gel regions were excised and de-stained 3 times in 50% methanol for 15 minutes each time. Then the gel regions were treated with 50 mM NH4HCO3 in 30% acetonitrile until completely de-stained. Finally, each gel sample was dehydrated with 100% acetonitrile. Dried gel samples were digested overnight with mass spectrometry grade trypsin (Trypsin Gold, Promega, Madison, WI, USA) at 5ng/μl in 50 mM NH4HCO3 digest buffer. After acidification with 10% formic acid, peptides were extracted with 5% formic acid/50% Acetonitrile (v/v) and concentrated to a small droplet using vacuum centrifugation. Desalting of peptides was done using hand packed SPE Empore C18 Extraction Disks as described (Rappsilber et al, 2007). Desalted peptides were again concentrated and reconstituted in 10 μL 0.1% formic acid in water. Aliquots of the peptides were analyzed by nano-liquid chromatography followed by tandem mass spectrometry (nano-LC-MS/MS) using an Easy nLC 1000 equipped with a self-packed 75 μm x 20 cm reverse phase column (ReproSil-Pur C18, 3 μm, Dr. Maisch GmbH, Germany) coupled online to a QExactive HF Orbitrap mass spectrometer via a Nanospray Flex source (all instruments from Thermo Fisher Scientific, Waltham, MA, USA). Analytical column temperature was maintained at 5O°C by a column oven (Sonation GmBH, Germany). Peptides were eluted with a 3-40% acetonitrile gradient over 110 min at a flow rate of 250 nL/min. The mass spectrometer was operated in data-independent-analysis mode and profile mode with survey scans acquired at a resolution of 120,000 (at m/z 200) over a scan range of 300-1650 m/z. Following the survey scans, 30 groups of precursors were selected for fragmentation with sliding isolation windows to include peptide m/z values ranging from 364 to 1370 Th. The default maximum charge state was set to 4 and resolution was set to 30,000. In MS/MS, the fixed first mass was set to 200 Da. The normalized collision energy was 27. The maximum injection times for the survey was set to 60 ms and the maximum injection time of MS/MS scans was set to auto. The ion target value for both scan modes was set to 3e6. Data were analyzed by the software MaxQuant (version 2.1.1.0), referred to as MaxDIA analysis type (Sinitcyn et al, 2021). To identify peptides, we used mouse in silico-generated spectral libraries that were created from the Uniprot mouse protein sequence database (downloaded on 06/18/2020; 21,989 entries). The library was obtained from the Max Planck Institute of Biochemistry Data share drive (MPIB Datashare https://datashare.biochem.mpg.de/s/qe1IqcKbz2j2Ruf?path=%2FDiscoveryLibraries). To determine the relative abundance of a protein across a set of samples, we used its label free quantitation (MaxLFQ) intensity calculated by MaxDIA (Sinitcyn et al, 2021), which normalizes the protein intensity by normalizing peptide intensities (minimum peptide ratio count=2) across samples. To determine the relative abundance of a protein complex, we summed LFQ intensities of all observed subunits.

### Turnover measurements with stable isotopes

In order to measure the turnover of proteins and lipids, flies were kept on media that contained either heavy lysine or heavy glucose. Flies were harvested after different incubation periods to determine the incorporation of heavy isotopes into proteins and lipids of indirect flight muscles by MS.

The fly food used to label Drosophila melanogaster was adopted from a previous publication (Sury et al, 2010). The medium consisted of 150 mg glucose and 30 mg yeast paste, which were mixed in a small amount of water and placed on a 0.25×0.25 inch^2^ square of filter paper. For the measurement of protein turnover, ^13^C_6_^15^N_2_-lysine-labeled yeast was mixed with unlabeled glucose. For the measurement of lipid turnover, unlabeled yeast was mixed with ^13^C_6_-glucose. Stable isotopes were obtained from Cambridge Isotope Lab. Inc. (Andover, MA 01810 USA). To prepare yeast, the lysine auxotrophic yeast strain SUB62/DF5 (MATalpha lys2-801 leu2-3/112 ura3-52 his3-delta200 trp1-1), obtained from the American Type Culture Collection (ATCC #200912), was grown on a medium containing 1.7 g/L yeast nitrogen base (without amino acids, without ammonium sulfate), 20 g/L D -glucose, 5 g/L ammonium sulfate, 200 mg/L adenine hemisulfate, 20 mg/L uracil, 100 mg/L tyrosine, 10 mg/L histidine, 60 mg/L leucine, 10 mg/L methionine, 60 mg/L phenylalanine, 40 mg/L tryptophan, 100 mg/L arginine, and 30 mg/L ^13^C_6_^15^N_2_-lysine or unlabeled lysine, respectively. LC-MS/MS analysis demonstrated that >99% of proteins were labeled when yeast cells were grown in the presence of heavy lysine.

To reduce variability, only male flies were studied. Flies were kept at 24°C in a moisturized chamber containing the glucose/yeast media specified above. Batches of 15 animals were collected per measurement. Flies were harvested after 2 and 4 days. Thoraces were dissected under the microscope and stored at −80°C until further use. Thoraces were homogenized in water and processed for protein and lipid analysis. In order to analyze proteins, the digestion was carried out with LysC instead of trypsin. The mass spectrometer was operated in data-dependent-analysis mode with survey scans acquired at a resolution of 70,000 (at m/z 200) over a scan range of 300-1750 m/z. Up to 10 of the most abundant precursors from the survey scan were selected with an isolation window of 1.6 Th for fragmentation by higher-energy collisional dissociation with a normalized collision energy of 27. The maximum injection times for the survey and MS/MS scans were 60 ms and the ion target value for both scan modes was set to 3e6. Data were analyzed by the software MaxQuant (version 1.5.5.1). We used the Andromeda search engine (Cox et al, 2011; Tyanova et al, 2016) to search the Uniprot Drosophila protein sequence database (downloaded on 01/09/2015; 3,214 entries). In order to analyze lipids, the lipidomics protocol described above was followed, except that spectra were acquired in centroid mode.

Principles of the kinetic analysis were previously described for proteins (Ross et al, 2021) and lipids (Schlame et al, 2020). Protein mass spectra were processed by the software MaxQuant (version 1.5.5.1) to yield heavy/light abundance ratios (*r*) for each identified protein. The ratio was converted into the relative isotopomer abundance (*r/1+r*) and divided by the relative abundance of heavy lysine in the free lysine pool of flight muscle (*p*) in order to yield the fractional synthesis (*q*), i.e. the proportion of newly synthesized protein in the total amount of protein present at time *t*:

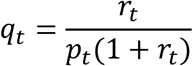

The relative isotopomer abundance was divided by *p* in order to correct for the presence of unlabeled lysine in the free lysine pool, which dilutes the abundance of heavy isotopes in newly synthesized peptides. The relative abundance of heavy lysine in the free lysine pool was determined by LC-MS/MS analysis on a metabolomics platform (Schlame et al, 2020). Turnover rate constants (*k*) were estimated from serial fractional syntheses measured at different time points (*t*) by non-linear regression to the equation:

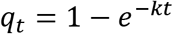

Half-life times were calculated as:

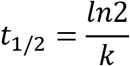

Lipid mass spectra were processed in the Qual Browser of Xcalibur (version 4.1.50). Lipid species with favorable signal-to-noise ratios were manually selected for analysis. In most lipids, we measured the abundance of the light isotopomer (monoisotopic peak) and the abundance of the heavy isotopomer (^13^C_3_ peak) and applied the same data processing algorithm as for proteins. In the lipid work-flow, *p* is the fraction of ^13^C_3_-glycerol-3-phosphate in the glycerol-3-phosphate pool of flight muscle, which was determined by the same method used for lysine. In case of PG and CL, several isotopomers were measured and the data were processed as described (Schlame et al, 2020).

## RESULTS

### Cardiomyocyte-restricted *Crls1*-KO causes dilated cardiomyopathy in infant mice

To determine whether CL affects cardiac development in mice, we inactivated *Crls1* in cardiomyocytes using *Cre-Lox* recombination with the *Myh6-Cre* driver. We compared *Myh6-Cre/+; Crls1^flox/flox^* (*Crls1*KO) mice to their +/+; *Crls1^flox/flox^* littermate controls. *Crls1*KO mice were born at the expected Mendelian ratio but had an average lifespan of only 16 days (**Figure 1A**). Early lethality could be attributed to heart failure. Although no cardiac defects were detectable at birth, *Crls1*KO mice developed severe dilated cardiomyopathy within 2 weeks as demonstrated by thinned ventricular walls (**Figure 1B**), increased diastolic dimension (**Figure 1C**), and reduced shortening fraction (**Figure 1D**). Thus, cardiomyocyte-restricted KO of *Crls1* had little effect on prenatal development but induced a lethal cardiomyopathy before the weaning age.

**Figure 1.**
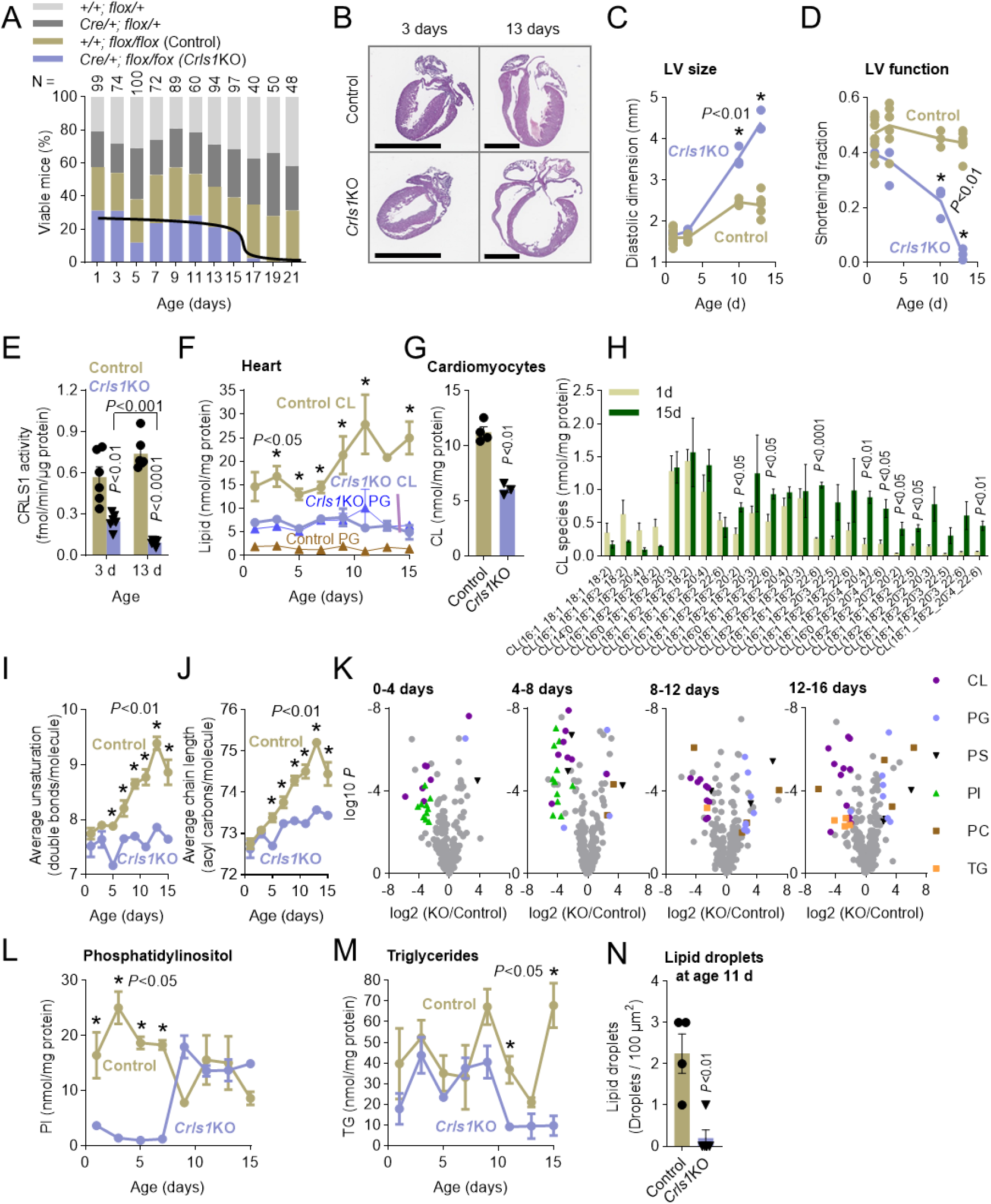
Survival, heart function, CRLS1 activity, and cardiac lipidome in *Crls1*KO and control mice. (A) Offspring were genotyped at different postnatal ages. (B) Sections of mouse hearts were stained with hematoxylin and eosin. Scale bars: 2 mm. (C-D) The left ventricle (LV) of mouse hearts was analyzed by echocardiography (N≥3). (E) CRLS1 activity was measured in heart (N=6). (F) CL and PG were quantified in heart (N=3). (G) CL was quantified in purified cardiomyocytes from 3-day old mice (N=4). (H) CL was analyzed in hearts of control mice at ages 1 day and 15 days (N=3). (I-J) CL was analyzed in hearts of *Crls1*KO and control mice at different ages (N=3). (K) Volcano plots compare cardiac lipids of *Crls1*KO (N=6) and control (N=6) mice at different ages. Species with significant differences between genotypes (4-fold higher or lower, *P*<0.01) are shown in color. (L-M) Lipids were quantified in hearts at different ages (N=3). (N) The density of lipid droplets was quantified in EM images of mouse hearts (N=4). Data are means with standard error. Comparisons were made by *t*-test.

To quantify the KO efficiency, we measured the gene product CRLS1. Because suitable antibodies for quantitative Western blotting of CRLS1 are not available, we developed an enzymatic assay for CRLS1 activity (**Figure S1**). *Crls1*KO reduced the enzymatic activity of CRLS1 in newborn mouse hearts by >50% and caused further decline in the activity as mice grew older (**Figure 1E**). Next, we followed the evolution of the lipidome in developing mouse hearts by liquid-chromatography/tandem-mass-spectrometry (LC-MS/MS) with internal standards. Close to 800 molecular species were identified, among them 94 molecular species of CL (**Data S1**). The analysis confirmed that, at birth, the concentration of CL was reduced by half while the concentration of the CRLS1 substrate phosphatidylglycerol (PG) was increased 3-fold. Importantly, control mice almost doubled the cardiac concentration of CL during the first two weeks of life (14.7±3.2 vs 23.7±6.5 nmol/mg protein; N=12; P<0.001) whereas *Crls1*KO mice maintained the concentration at the natal level (7.0±1.0 vs 6.3±2.2 nmol/mg protein; N=12; P=0.32) (**Figure 1F**). To determine whether the residual CL in *Crls1*KO hearts was contributed by cardiomyocytes, rather than fibroblasts or endothelial cells, we purified cardiomyocytes from 3-day old mouse hearts and determined their lipid composition. Indeed, *Crls1*KO cardiomyocytes still contained about half the CL concentration of normal cardiomyocytes (**Figure 1G**). Thus, KO of *Crls1* in cardiomyocytes caused substantial loss of CRLS1 and CL, increase in PG, and failure to boost the CL concentration during postnatal development.

In control mice, postnatal development induced drastic changes in the molecular species composition of CL because only long-chain and highly unsaturated species increased their concentration whereas shorter and more saturated species declined or did not change (**Figure 1H**). This led to a gradual rise in the average unsaturation and the length of the acyl chains linked to CL. However, in *Crls1*KO hearts, the unsaturation of CL remained unchanged during postnatal development (**Figure 1I**) and the acyl chain length increased only slightly (**Figure 1J**). We concluded that postnatal development was associated with CL remodeling. In *Crls1*KO mice, remodeling of the residual CL was inhibited even though the KO was not directed at the remodeling enzyme tafazzin.

In order to identify other alterations in the lipidome, we compared *Crls1*KO and controls in volcano plots. Apart from the primary effects on CL and PG, *Crls1*KO altered molecular species of phosphatidylinositol (PI), triglyceride (TG), and to a lesser extent, phosphatidylserine (PS) and phosphatidylcholine (PC). With increasing postnatal age, the effects on CL, PG, and TG became stronger whereas the effect on PI disappeared (**Figure 1K**). This was because *Crls1*KO mice were born with an abnormally low PI concentration, which corrected itself during postnatal development (**Figure 1L**). In contrast, the concentration of TG was normal at birth but dropped suddenly a few days before death in *Crls1*KO mice (**Figure 1M**). At the same time, we observed a loss of lipid droplets (**Figure 1N**), suggesting the exhaustion of metabolic fuel stores in the failing *Crls1*KO hearts.

In summary, cardiomyocyte-restricted KO of *Crls1* substantially reduced CRLS1 activity and CL concentration at birth and prevented the rise of CL and the remodeling of its acyl chains during postnatal development. *Crls1*KO mice were born with near normal cardiac function but quickly developed severe dilated cardiomyopathy, which led to their death at the age of 2 weeks. The data suggest that postnatal cardiac development is critically dependent on a progressive rise in CL and on the remodeling of its acyl chains.

### CL deficiency prevents the accumulation of OXPHOS proteins during cardiac maturation

To compare the postnatal maturation of cardiomyocytes in *Crls1*KO mice and controls, we determined the protein composition of developing mouse hearts at multiple time points. Using label-free relative quantitative proteomics, we identified 5,002 proteins, which belonged to the nucleus (31%), the mitochondria (15%), the plasma membrane (15%), the cytosol (11%), the endoplasmic reticulum (8%), the myofibrils (8%), and other compartments (**Data S2**). First, we analyzed the correlation between protein abundances and age in control hearts, which demonstrated an asymmetric effect of maturation on the protein composition. While 3,279 proteins (66%) decreased during maturation (Pearson correlation coefficient *r* < −0.5) only 64 proteins (1.3%) increased during maturation (*r* > +0.5). Of those 64 proteins, 24 (38%) belonged to mitochondria. *Crls1*KO did not affect the number of proteins with negative correlation but reduced the number of proteins with positive correlation. Only 7 out of 34 proteins with positive correlation (21%) belonged to mitochondria in *Crls1*KO hearts (**Figure 2A**). Thus, KO of *Crls1* inhibited the rise of mitochondrial proteins during cardiomyocyte maturation.

**Figure 2.**
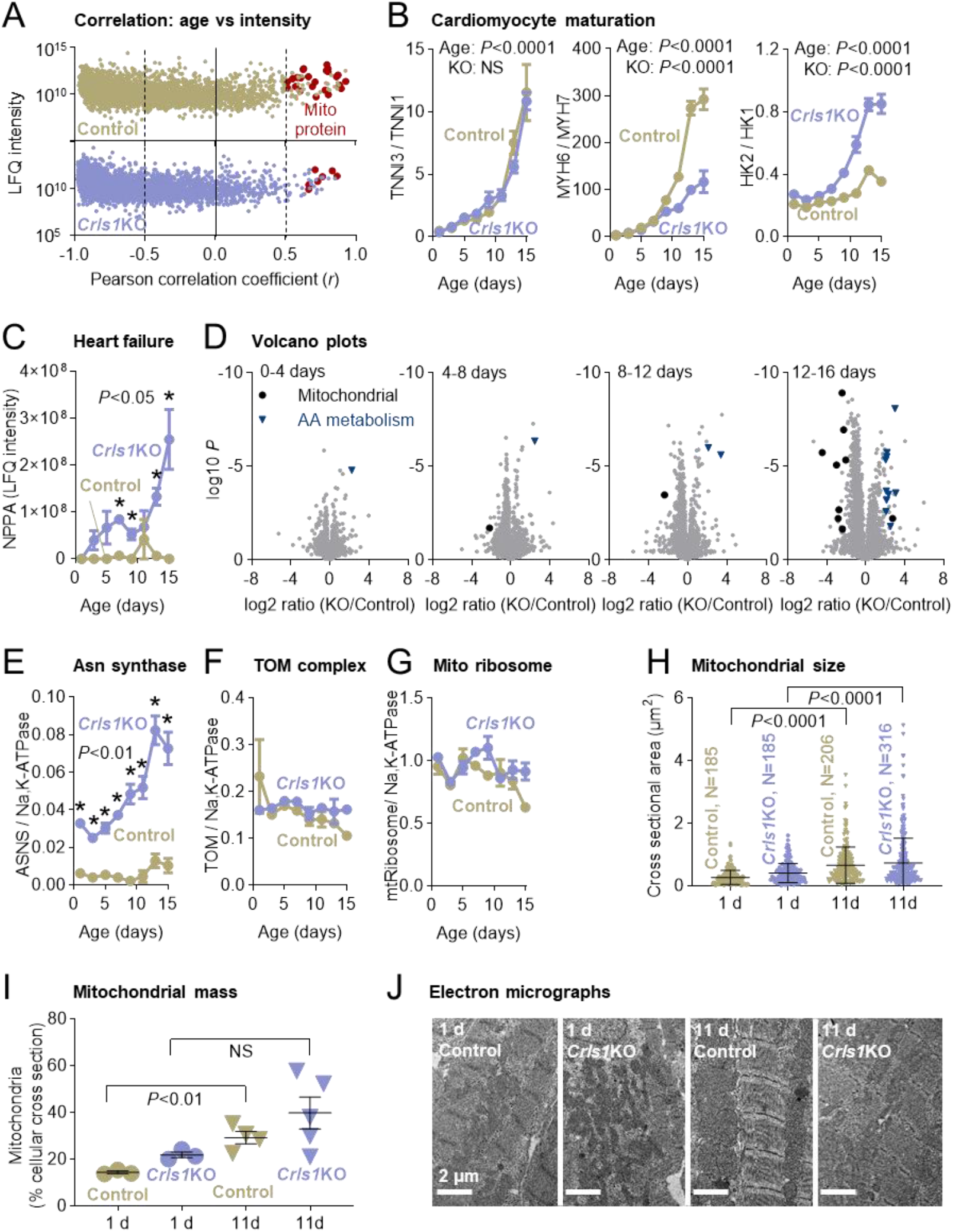
Maturation of cardiomyocytes in *Crls1*KO and control mice. (A-G) Heart samples (N=3) were collected at different ages and analyzed by label-free relative quantitative proteomics. Correlations between protein abundances and age were determined by calculating Pearson correlation coefficients (*r*). Mitochondrial proteins are marked in the group with significant positive correlation (*r* > 0.5) (A). Data are means with standard error (B, C, E, F). Comparisons between *Crls1*KOs and controls were made by two-way-ANOVA (B), *t*-tests (C, E-G), and volcano plots (D). Volcano plots were created of 4 different age groups. Proteins that were significantly different between genotypes (4-fold higher or lower, *P*<0.01) are marked if they are localized in mitochondria or are involved in amino acid (AA) metabolism. Asn, asparagine; NPPA, atrial natriuretic peptide; NS, not significant; TOM, translocase of the outer membrane. (H-J) Heart samples were analyzed by quantitative EM. Data are means with standard deviation (H) or standard error (I). Comparisons were made by *t*-tests. N, number of mitochondrial cross sections; NS, not significant.

Next, we analyzed specific indicators of cardiac maturation by measuring ratios of adult versus fetal isoforms of 3 proteins that are known to undergo isoform switching during maturation (Guo & Pu, 2020). They included troponin I (TNNI), myosin heavy chain (MYH), and hexokinase (HK). Isoform switching is designed to improve the efficiency of contraction and to convert the energy metabolism from glycolysis to OXPHOS. As expected, the adult/fetal isoform ratios increased steadily in control hearts over the first two weeks of life. *Crls1*KO did not alter the progression of the TNNI3/TNNI1 ratio but slowed the rise of the MYH6/MYH7 ratio and accelerated the rise of HK2/HK1 (**Figure 2B**). Furthermore, *Crls1*KO caused progressive rise in the atrial natriuretic peptide (NPPA), a marker of heart failure (Goetze et al, 2020) that was virtually absent from controls. The increase in the atrial natriuretic peptide occurred parallel to the decline in cardiac function (**Figure 2C**). The data demonstrate that *Crls1*KO altered the characteristics of cardiomyocyte maturation and caused heart failure in newborn mice.

To pinpoint the age at which changes in the proteome occur, we created volcano plots of data collected at sequential time points. Differences between *Crls1*KO and control became gradually larger during postnatal development. Proteins that decreased in *Crls1*KO belonged mostly to the mitochondrial compartment while proteins that increased in *Crls1*KO were mostly active in amino acid metabolism (**Figure 2D**). For instance, *Crls1*KO substantially increased the relative abundance of asparagine synthase (**Figure 2E**). Among the mitochondrial proteins suppressed by *Crls1*KO were many OXPHOS subunits. However, proteins involved in mitochondrial biogenesis, such as the translocase of the outer membrane (TOM) (**Figure 2F**) and mitochondrial ribosomes (**Figure 2G**), were not affected. Quantitative electron microscopy (EM) demonstrated an increase in mitochondrial size (**Figure 2H**), total mass (**Figure 2I**), and alignment with myofibrils (**Figure 2J**) during postnatal development but no effect of *Crls1*KO on these parameters. We concluded that *Crls1*KO mice were born with a normal cardiac proteome but developed aberrancies during cardiomyocyte maturation. Although these aberrancies affected mostly mitochondrial proteins, they did not prevent the expansion of the mitochondrial compartment during maturation.

Since *Crls1*KO altered the abundance of some mitochondrial proteins but not the mitochondrial mass, we asked whether *Crls1*KO changed the internal composition and the internal structure of mitochondria. First, we measured crista densities by EM, which revealed an increase in the number of cristae during development (**Figure 3A**). *Crls1*KO did not affect the crista density at birth but diminished its subsequent rise (**Figure 3B**). We noticed that a portion of *Crls1*KO mitochondria (17±8% in 15 images) contained crista-free zones, which contributed to the paucity of cristae (**Figure 3C**). Thus, *Crls1*KO caused a slight inhibition in the rise of crista density during cardiac development.

**Figure 3.**
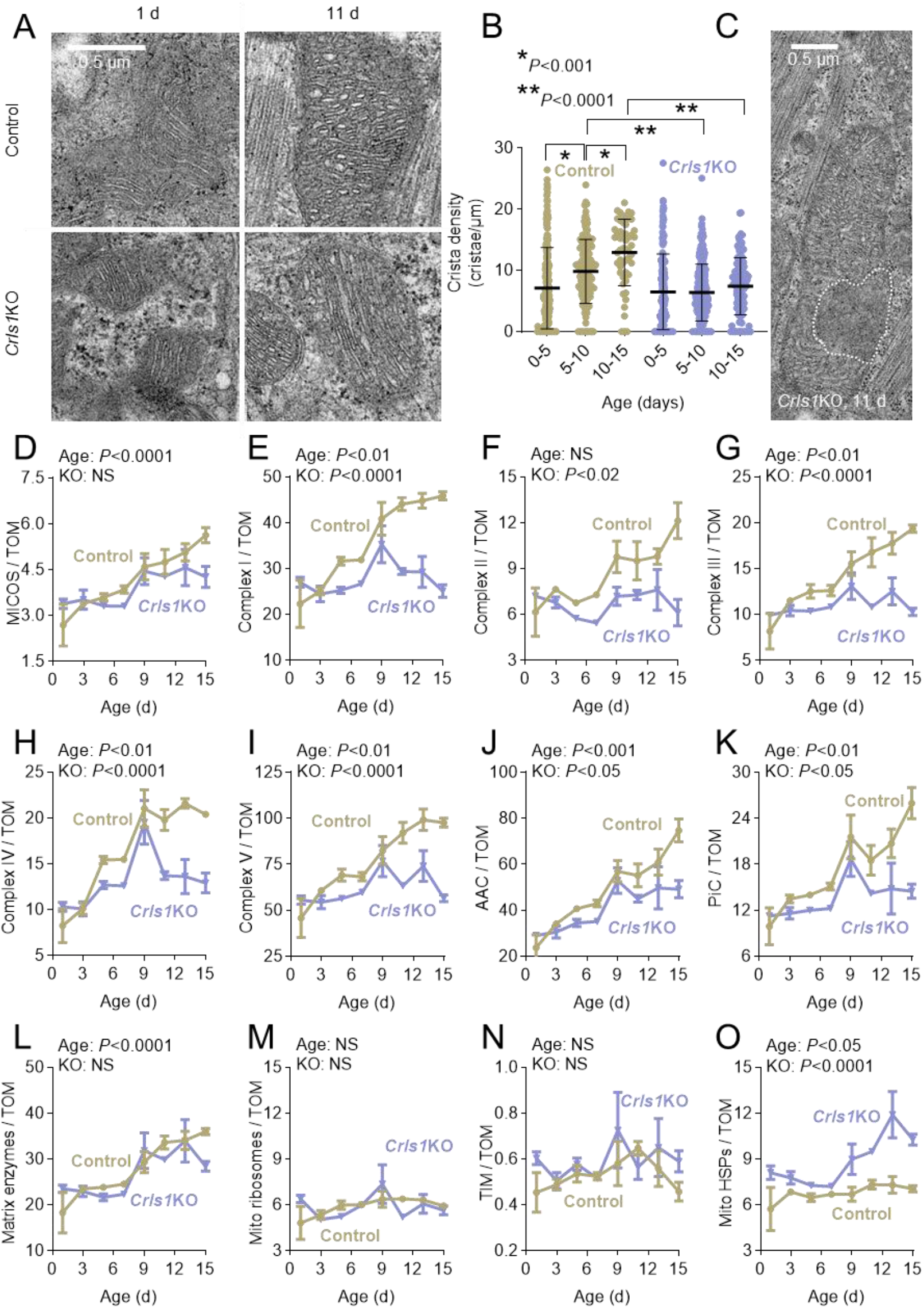
Internal structure and protein composition of cardiac mitochondria in *Crls1*KO and control mice. (A-C) Heart samples were analyzed by quantitative EM. The dotted line in panel C marks a crista-free zone. The graph shows means with standard deviation of >100 mitochondrial cross sections in each group. Comparisons were made by *t*-test. (D-O) Heart samples were analyzed by label-free relative quantitative proteomics in order to calculate ratios of the abundance of protein complexes or protein groups over the abundance of the translocase of the outer membrane (TOM). Data are means with standard error (N=3). Effects of age and *Crls1*-knockout (KO) were analyzed by two-way-ANOVA. AAC, ADP/ATP carriers; MICOS, mitochondrial contact site and cristae organizing system; PiC, phosphate carrier; TIM, translocase of the inner membrane; HSPs, heat shock proteins; NS, not significant.

Second, we determined the intramitochondrial concentration of protein complexes. To this end we summed the label-free-quantitation (LFQ) intensities of individual subunits and calculated the ratio of this sum to the LFQ intensity of the TOM complex. TOM was used as reference because, as an outer membrane protein, its abundance correlates with the outer surface area and therefore with the volume of mitochondria. We found that the mitochondrial concentration of the crista junction complex MICOS increased during cardiac development, which was consistent with the increase in crista density by EM. However, *Crls1*KO did not have a significant effect on the MICOS concentration (**Figure 3D**). In contrast, the mitochondrial concentration of all 5 OXPHOS complexes steadily increased during cardiomyocyte maturation only in normal but not in *Crls1*KO mice (**Figure 3E-I**). The same was true for the ADP/ATP carriers (**Figure 3J**) and the phosphate carrier (**Figure 3K**). In comparison, *Crls1*KO did not have any effect on the mitochondrial concentration of matrix proteins, such as matrix enzymes (**Figure 3L**) and mitochondrial ribosomes (**Figure 3M**), and also not on the translocase of the inner membrane (TIM), which is located in the inner boundary membrane (**Figure 3N**). Finally, *Crls1*KO increased the concentration of heat shock proteins in the mitochondrial matrix, mostly in the second week of life, suggesting increased unfolding of proteins in the mutant mitochondria (**Figure 3O**). We concluded that *Crls1*KO prevented specifically the rise in the concentration of crista proteins, such as OXPHOS complexes and carriers, during cardiomyocyte maturation.

In summary, *Crls1*KO mice were born with normal heart mitochondria but failed to increase the intramitochondrial concentration of OXPHOS proteins and carriers during cardiomyocyte maturation. The KO effect was specific for proteins of the crista membrane and did not apply to proteins of the matrix or the inner boundary membrane. Abnormalities of the proteome were progressive and ultimately caused severe stress as indicated by markers of heart failure and heat shock response.

### CL prolongs the lifetime of OXPHOS proteins in Drosophila flight muscle

Lack of OXPHOS complexes can be caused by reduced synthesis of OXPHOS proteins or by increased degradation. The underlying mechanism may affect specific subunits, whole complexes, or the entire crista membrane including proteins and lipids. To distinguish between these possibilities, we measured the turnover rate of proteins and lipids with stable isotopes. These experiments were carried out in Drosophila where parallel labeling of proteins and lipids is technically feasible and CL-deficient mutants are available. We limited our analysis to indirect flight muscles because, like mammalian hearts, they contain crista-rich mitochondria and perform energy-intensive contractions that are sensitive to *Crls1* deletion (Acehan et al, 2011).

To measure the turnover of proteins and lipids, we incubated flies on media that contained either ^13^C_6_^15^N_2_-lysine (protein synthesis) or ^13^C_6_-glucose (lipid synthesis) but had otherwise identical compositions. Muscles were harvested by thorax excision at different time points and analyzed by LC-MS/MS for proteomics or lipidomics, respectively. After digestion of proteins by LysC, peptides were recovered as light (m/z=M) and heavy (m/z=M+8) isotopomers in MS1 spectra. Most lipids were also recovered as light (M) and heavy (M+3) isotopomers, except for PG (M, M+3, M+6) and CL (M, M+3, M+6, M+9), which formed multiple isotopomers (**Figure 4A**) (Schlame et al, 2020). The heavy/light intensity ratios of peptides and lipids increased over time as a result of the incorporation of heavy isotopes (**Figure 4B**). Since adult flies do not accumulate body mass over time, the incorporation of heavy isotopes was the result of the turnover of proteins and lipids. We calculated fractional syntheses from heavy/light intensity ratios and estimated the corresponding rate constants of turnover by non-linear regression analysis (**Figure 4C**). Rate constants were transformed into half-life times based on the assumption of steady state.

**Figure 4.**
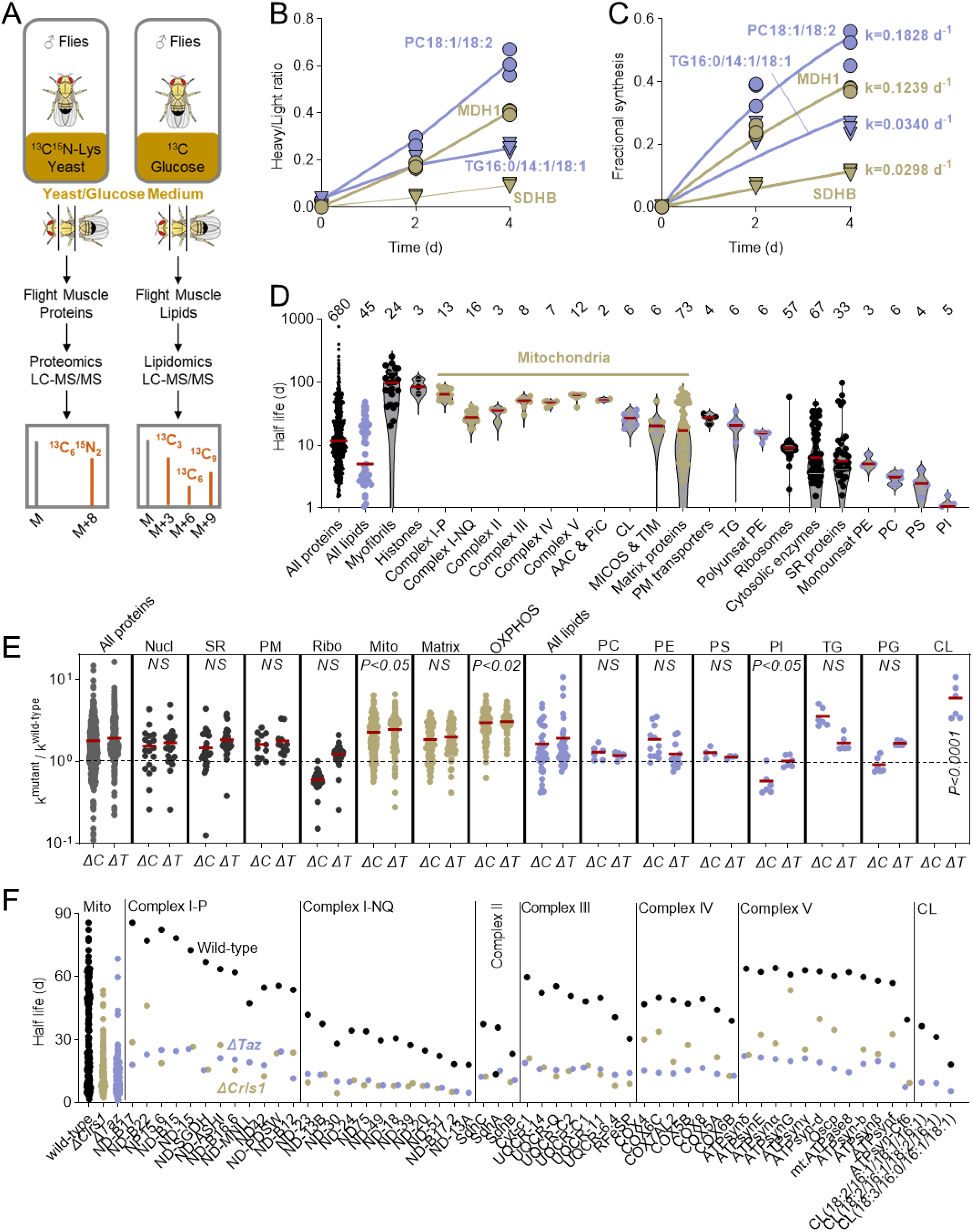
CL deficiency shortens the lifetime of OXPHOS subunits in Drosophila flight muscles. (A-C) Male flies were incubated on yeast/glucose containing either heavy lysine or heavy glucose. Flight muscles were harvested and analyzed by LC-MS/MS to determine isotopomer patterns of peptides or lipids (3-4 replicas per time point, 15 flies per replica). Fractional syntheses were calculated from heavy/light intensity ratios. Turnover rate constants (k) were estimated by non-linear regression analysis. (D) Half-lives were calculated from rate constants. Columns show the half-lives of individual proteins and lipids and their median (red line). The sample size of each category is shown on top. (E) Data are mutant/wild-type ratios of the turnover rate constants of proteins and lipids. Red lines indicate means. Groups were compared by *t*-test against the ratios of the entire proteome or lipidome, respectively. ΔC, *ΔCrls1;* ΔT, *ΔTaz*. (F) Data are half-lives of OXPHOS subunits and CL species in wild-type, *ΔCrls1*, and *ΔTaz*. In all complexes and CL, there was a significant difference between the wild-type and the mutants (*P*<0.05; t-test). The half-lives of all mitochondrial proteins are shown for comparison. AAC, ADP-ATP carrier; PiC, phosphate carrier; MICOS, mitochondrial contact site and cristae organizing system; TIM, translocase of the inner membrane; PM, plasma membrane; TG, triglyceride; PE, phosphatidylethanolamine; SR, sarcoplasmic reticulum; PC, phosphatidylcholine; PS, phosphatidylserine; PI, phosphatidylinositol.

We analyzed the turnover of 680 proteins and 48 lipids in Drosophila flight muscle (**Data S3**). In wild-type, the majority of proteins and lipids had half-lives of 1-100 days. The average lifetime of lipids was shorter than that of proteins. OXPHOS complexes belonged to the longest-lived proteins together with histones and myofibrils. Within the OXPHOS system, the P module of complex I had the longest lifespan and the NQ module of complex I had the shortest. The long lifetimes of OXPHOS proteins (Bomba-Warczak et al, 2021; Fornasiero et al, 2018; Kim et al, 2012; Price et al, 2010) and the discrepancy between the P and NQ-modules of complex I (Krishna et al, 2021; Szczepanowska et al, 2020) have already been described in mammals. However, not all mitochondrial proteins were as stable as OXPHOS complexes. In particular, the mitochondrial matrix contained proteins with a much shorter lifespan. Among lipids, lifetimes were class-specific except for PE where mono-unsaturated species turned over faster than polyunsaturated species. The lipid with the longest lifetime was CL and the lipid with the shortest was PI (**Figure 4D**). Together the data demonstrate a remarkable longevity of both CL and OXPHOS proteins, suggesting that membrane domains harboring the OXPHOS system are largely inert to degradation.

We measured protein and lipid turnover in two CL-deficient mutants, one with inactivated CL synthase (*ΔCrls1*, resulting in complete loss of CL) and one with inactivated tafazzin (*ΔTaz*, resulting in low concentration and abnormal composition of CL). Rate constants of individual proteins and lipids in the mutants were divided by the corresponding rate constant in the wild-type. In either mutant, CL deficiency decreased the turnover of some proteins (k^mutant^/k^wild-type^<1) but increased the turnover of others (k^mutant^/k^wild-type^>1). The average k^mutant^/k^wild-type^ ratio was 1.7 in *ΔCrls1* and 1.9 in *ΔTaz*,indicating a slight increase in the overall protein turnover as the result of CL deficiency. However, the turnover of OXPHOS proteins stood out in that it was about 3-fold higher in the mutants than in the wild-type. In contrast, proteins of the mitochondrial matrix and proteins of other cellular compartments did not show a large increase in turnover. Among phospholipids, only CL experienced increased turnover in *ΔTaz* whereas PC, PE, PS, and PG remained unaffected. Furthermore, *ΔCrls1*, but not *ΔTaz*, decreased the turnover of PI and increased the turnover of TG (**Figure 4E**). We speculate that the increase in TG turnover results from inefficient energy metabolism, which requires increased TG oxidation.

Thus, our data indicate that CL deficiency increased the turnover of OXPHOS proteins and, in the case of *ΔTaz* where some CL remains, the turnover of CL itself. Indeed, CL deficiency, caused either by *ΔCrls1* or *ΔTaz*, reduced the half-life of every OXPHOS subunit identified in the proteome. Proteins with the longest lifetime, such as the subunits of the P module of complex I or the subunits of complex V, were most affected. As a result, all OXPHOS proteins and CL species assumed a similar half-life in the mutants, which was much shorter than their normal half-life (**Figure 4F**). Interestingly, the more saturated a CL species was the less stable it became (**Figure S2**). This observation is consistent with the idea that saturated CLs are less suited to interact with OXPHOS proteins (Xu et al, 2021).

In summary, we have shown that OXPHOS complexes are among the most stable proteins of Drosophila flight muscles. CL is essential for their long lifetime and has itself a long half-life, suggesting that CL and OXPHOS complexes are co-preserved and that their interaction protects them from degradation. Either the loss of CL remodeling (*ΔTaz*) or the lack of CL (*ΔCrls1*), expose OXPHOS proteins to increased turnover. The fact that not only OXPHOS proteins but also CL had a shortened half-life in *ΔTaz*, suggests that once the CL-OXPHOS interaction is weakened, lipids and proteins of cristae are co-degraded.

## DISCUSSION

We have created a new murine model of neonatal cardiomyopathy by inactivating the CL-synthesizing enzyme *Crls1* in cardiomyocytes. These mice deteriorated quickly during the maturation period of the neonatal heart but lived long enough to study the effects of CL deficiency. Our data reveal the critical importance of CL for the accumulation of OXPHOS complexes in mitochondria during neonatal development. Remarkably, CL deficiency did not affect mitochondrial proteins outside the cristae nor did it affect mitochondrial size or mass. Instead, it prevented the cumulative rise in the concentration of OXPHOS proteins and carriers within mitochondria, which we observed during the maturation of normal mice. As a result, CL deficiency interfered with cardiac maturation, the process that transforms neonatal into adult cardiomyocytes (Guo & Pu, 2020), and thus prevented the creation of an efficiently contracting heart. Separate experiments in Drosophila flight muscle demonstrated that CL deficiency exposes OXPHOS complexes to increased degradation, which if true in cardiomyocytes, explains the failure to accumulate OXPHOS proteins in *Crls1*KO hearts. While high expenses and logistical challenges prevented us from replicating these experiments in mice, the Drosophila data confirm the well-documented longevity of OXPHOS proteins in different cells (Bomba-Warczak et al, 2021; Fornasiero et al, 2018; Kim et al, 2012; Krishna et al, 2021; Morgenstern et al, 2021; Price et al, 2010). By showing that CL is essential for the longevity of OXPHOS proteins we discovered perhaps one of the most important functions of CL.

The present data support and expand a mechanistic model of CL action, which asserts that CLs with long and unsaturated chains promote the accumulation of OXPHOS complexes because they reduce stress that arises from protein crowding in the lipid bilayer of mitochondrial cristae (Xu et al, 2019). We have shown that protein crowding triggers the remodeling of fatty acids attached to CL and that either the lack of CL or the lack of CL remodeling prevents protein crowding in mitochondrial membranes (Xu et al, 2021). Here we show (i) that CL is necessary to accumulate OXPHOS complexes and carriers in cardiomyocyte mitochondria during neonatal development, (ii) that the accumulation of these proteins is associated with an increase in chain length and unsaturation of CL species, and (iii) that a drop in the CL concentration inhibits both protein accumulation and CL remodeling. These data are consistent with the idea that CL is necessary to allow protein crowding (OXPHOS accumulation) while protein crowding triggers CL remodeling. This explains why CL remodeling is inhibited in *Crls1*KO mice. Recent cryo-EM data, demonstrating a transient attachment of tafazzin to nascent complex I, also support the notion that CL remodeling is involved in the assembly of the OXPHOS system (Schiller et al, 2022).

To comprehend the effects of CL, one has to bear in mind its strong but non-specific affinity to proteins, which seems to result from the unique combination of a small but rigid head group and a large but flexible hydrophobic moiety (Lewis & McElhaney, 2009). The rigid head group exposes the phosphates to ionic interactions with amino acids, while the flexible acyl groups confer adaptability to diverse protein surfaces. Due to its strong interaction with diverse proteins, CL increases the cohesion within lipid-protein complexes. This is evident from the stabilizing effect of CL on supercomplexes (Pfeiffer et al, 2003; Zhang et al, 2002), from the prolonged residence time of CL on protein surfaces (Duncan et al, 2016), and from the resistance of CL against detergent extraction (Beyer & Klingenberg, 1985; Zhou et al, 2011). High cohesion provides a plausible explanation for the long half-life of CL-protein complexes because cohesion limits diffusion and molecular motion and therefore accessibility to hydrolyzing enzymes. The same mechanism may be invoked to explain the slow mixing of OXPHOS complexes upon fusion of mitochondria (Wilkens et al, 2013). Thus, CL helps to form compact crista domains that are shielded from degradation and membrane remodeling. Presumably these domains form the structural and functional backbone of the ATP-generating system in cardiac mitochondria, which explains the significance of CL for the heart.

In summary, by deleting *Crls1* from cardiomyocytes, we have established a mouse model of progressive heart failure and early death that mimics clinical aspects of inborn cardiomyopathies. Mice were born with nearly normal cardiac function and a nearly normal cardiac proteome but deteriorated during infancy. Our data reveal that the postnatal development of cardiomyocytes requires CL for the maturation of mitochondrial cristae, a process that entails the accumulation of OXPHOS proteins, and that a key mechanism of CL is to protect OXPHOS proteins from degradation.

## ACKNOWLEDGEMENT & SOURCES of FUNDING

This work was supported in part by the National Institutes of Health (grant R01 GM115593 to MS, shared instrumentation S10 grants RR027990 and OD023659, and core center P30 grant NS050276 to TAN) and The Seventh District Foundation (to CKLP). We also acknowledge the NYU Langone Health Microscopy Laboratory for consultation and assistance with transmission electron microscopy. This shared resource was partially supported by the Cancer Center Support Grant P30CA016087 at the Laura and Isaac Perlmutter Cancer Center.

## AUTHOR CONTRIBUTIONS

M.R., Y.X., C.K.L.P., H.E., and M.S. conducted experiments. M.R., C.K.L.P., H.E., T.A.N., and M.S. analyzed data. C.K.L.P., T.A.N., and M.S. acquired funding. M.R. and M.S. jointly developed the ideas and underlying concepts of the study. M.S. wrote the original draft of the paper and coordinated research activities. All authors edited the manuscript.

## LEGEND TO THE FIGURES

**Figure S1.**
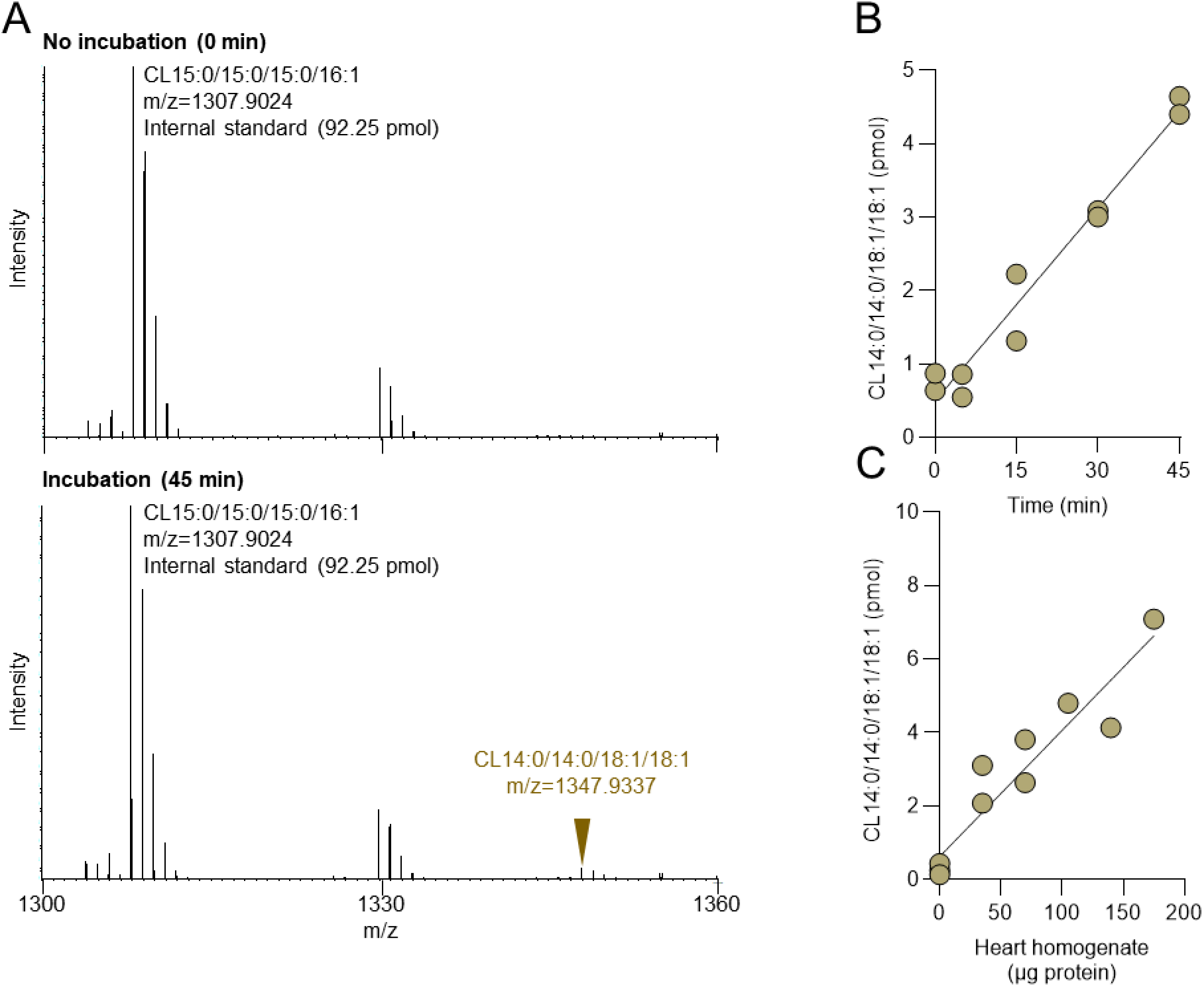
Assay of cardiolipin synthase (CRLS1) activity in heart homogenate. Mouse heart homogenate was incubated at 37°C with dimyristoyl-phosphatidylglycerol (PG14:0/14:0) and CDP-dioleoylglycerol (CDPDG18:1/18:1) at pH 8 in the presence of Co^2+^. The formation of CL14:0/14:0/18:1/18:1 was measured by LC-MS with internal standards. (A) CL14:0/14:0/18:1/18:1 is not an endogenous species of mouse heart but becomes detectable upon incubation with CRLS1 substrates. The panel shows sub-spectra (m/z 1300-1360 Th, retention time 12-15 min) that contain the reaction product CL14:0/14:0/18:1/18:1 and the internal standard CL 15:0/15:0/15:0/16:1. (B) CL14:0/14:0/18:1/18:1 increases at a constant rate during incubation. Assay mixtures contained 10 mM Tris (pH 8), 5 mM CoCl2, 0.1 mM PG14:0/14:0, 0.1 mM CDPDG18:1/18:1, and 70 μg heart protein in a total volume of 100 μL. Reactions were stopped after different time intervals. Lipids were extracted and CL was quantified by LC-MS. (C) Formation of CL14:0/14:0/18:1/18:1 is linearly dependent on the amount of heart homogenate. Assay mixtures contained 10 mM Tris (pH 8), 5 mM CoCl2, 0.1 mM PG14:0/14:0, 0.1 mM CDPDG18:1/18:1, and different amounts of heart homogenate in a total volume of 100 μL. Reactions were stopped after 30 minutes. Lipids were extracted and CL was quantified by LC-MS.

**Figure S2.**
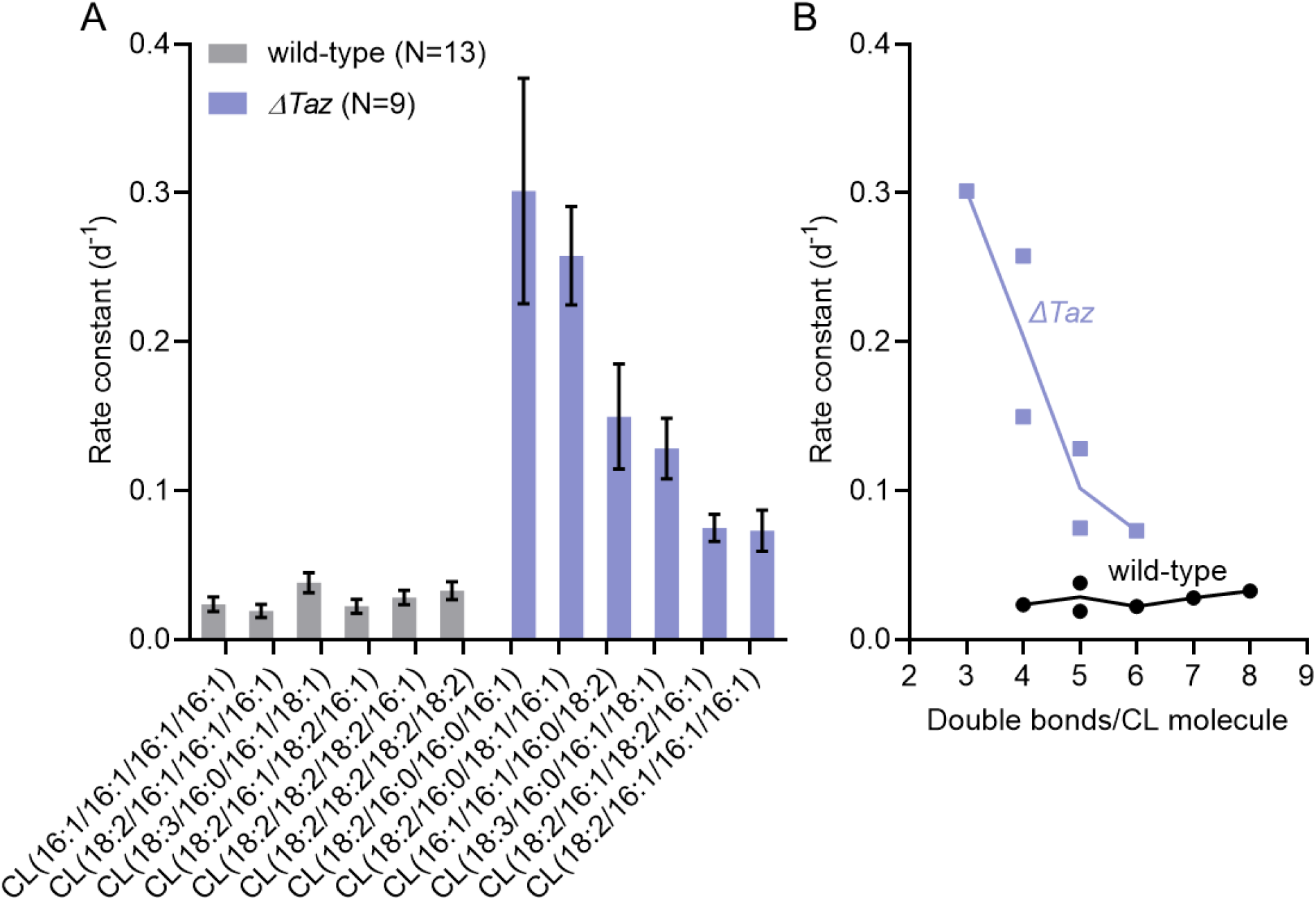
Stability of CL species in *ΔTaz* flight muscle depends on the number of double bonds. Wild-type and *ΔTaz* flies were incubated on a medium containing ^13^C_6_-glucose. The fractional syntheses of CL species were measured by LC-MS/MS in flight muscle. Rate constants were estimated by non-linear regression to the time dependence of fractional syntheses. (A) Data are rate constants of different CL species with the 95% confidence interval. N is the number of data points used for regression analysis. (B) The graph shows the dependence of rate constants on the number of double bonds per CL molecule.

